# Pathogenesis and host response to a novel Tacaribe virus isolate in experimentally-infected Jamaican fruit bats

**DOI:** 10.64898/2026.06.12.731631

**Authors:** Phillida A Charley, Luke Namesnik, Paul S. Soma, Ashley B. Reers, Clara Reasoner, Shijun Zhan, Bradly Burke, Liliana M Dávalos, Jan Felix Drexler, Allison C. Vilander, Rushika Perera, Hannah K Frank, Corey L. Campbell, Tony Schountz

**Affiliations:** Department of Microbiology, Immunology and Pathology, College of Veterinary Medicine and Biomedical Sciences, Colorado State University, Fort Collins, CO, USA; Department of Ecology and Evolutionary Biology, Tulane University, New Orleans, LA, USA; Department of Ecology and Evolution, Consortium for Inter-Disciplinary Environmental Research and Institute for Advanced Computational Science, Stony Brook University, Stony Brook, NY, USA; Institute of Virology, Charité, Universitätsmedizin Berlin, Germany

**Author notes:** Corresponding author; tony. These authors contributed equally to this project (Phillida A Charley, Luke Namesnik).

## Abstract

Tacaribe virus (TCRV) was the first arenavirus discovered in the New World and was isolated from *Artibeus* bats in Trinidad and Tobago in the 1950s. One isolate, TRVL-11573, remains but it was passaged by intracranial inoculation of newborn mice 22 times that likely changed its biology. This isolate has been extensively used for arenavirus research, including our previous work that showed it can cause fatal neurological disease in Jamaican fruit bats (*Artibeus jamaicensis*). Another divergent TCRV, DOM2014, was recently identified in transcriptome data from a Jamaican fruit bat captured in the Dominican Republic that contained TCRV genome. A kidney fragment homogenate from this bat was inoculated into Jamaican fruit bats and all became infected with signs of mild liver disease. Experimental challenge of Jamaican fruit bats with DOM2014 led to nonfatal infection that persisted through the end of the study on day 21 and with contact transmission to naive bats. Histopathology, immunohistochemistry and serum chemistry confirmed infection and mild liver disease, but none of the bats produced neutralizing antibodies. B cell receptor transcripts suggested limited somatic hypermutation that could explain the lack of detectable neutralizing antibodies. Transcriptome profiling of livers and spleens showed signatures of a typical innate antiviral response; however, evidence of adaptive immune suppression was also present. Similarly, liver transcriptome analysis showed signatures of an expected innate antiviral response and metabolic dysfunction. The isolation of TCRV DOM2014 provides a relevant model for the study of a bat reservoir host, and which may challenge the extensive work previously conducted with TRVL-11573.

**IMPORTANCE:** Several New World arenaviruses cause disease in humans and many are BSL-4 agents. TCRV strain TRVL-11573 has been used since the 1950s to study arenavirus biology; however, because it was intracranially passaged in newborn mice and Vero cells, it likely accumulated mutations that changed its biology. This assertion has been reinforced in recent years with discovery of divergent TCRV sequences in wild *Artibeus* bats that are substantially different than TRVL-11573, thus the prototype strain is unlikely to represent wildtype TCRV. The isolation of a new TCRV strain that has retained its genome fidelity allows a better understanding of TCRV’s biology and pathogenic potential. The availability of pathogen-free Jamaican fruit bats and cell lines that are permissive for TCRV DOM2014 will also help retain the biological features of the virus. Collectively, this is among the most tractable bat reservoir host models developed and provides a system for dissection of how bats host viruses. Moreover, it may be a suitable model for the study of therapeutics and vaccines for New World arenaviruses.

## INTRODUCTION

Tacaribe virus (TCRV) was first isolated in 1956 from a great fruit-eating bat (*Artibeus lituratus*) near Port-of-Spain, Republic of Trinidad and Tobago (1). Isolates were made from ten other *Artibeus* bats and one mosquito pool during the following two years by investigators at the Trinidad Regional Virus Laboratory (TRVL); however, only the original isolate, strain TRVL-11573, remains. The virus was intracranially passaged in newborn mice 22 times in the 1950s, likely altering its biology and, perhaps, enhancing neurotropism. One other isolate was made from a Lone Star tick (*Amblyomma americanum*) pool collected in central Florida in 2012 that surprisingly shared >99% nucleotide identity with TRVL-11573 (2). The natural history of this isolate is unclear because of its features that are inconsistent with arenavirus biology; arenaviruses are not known to be transmitted by ticks and the putative *Artibeus* bat reservoirs or other fruit bats are not found in mainland Florida. Divergent TCRV genomes also were recently characterized in four great fruit-eating bats and one flat-faced fruit bat (*A. planirostris*) sampled in Brazil but no isolates were made (3). TCRV sequences have also been detected in insectivorous velvety free-tailed bats (*Molossus molossus*) in Brazil, representing a potential spillover infection, since the detected virus was genetically closely related to *Artibeus*-borne TCRV (4).

TCRV is a clade B New World (NW) arenavirus that includes several viruses with high infection fatality rates that are hosted by rodents (5), including Machupo virus (MACV), Junìn virus (JUNV), and Guanarito virus (GTOV) (6). Only one human infection with TCRV has been documented, a laboratory-acquired infection with TRVL-11573 that caused febrile illness, recovery and high titer seroconversion (7). The viruses use transferrin receptor-1 (Tfr1) for cellular entry (8) that is found on many cell types. However, TRVL-11573 cannot use human Tfr1 whereas the others can, which may account for their substantial pathogenesis in humans (6). Because of its passage history, TCRV TRVL-11573 may be attenuated for human infection.

We previously determined that high-dose TRVL-11573 experimental infection of Jamaican fruit bats from a pathogen-free, captive breeding colony led to fatal neurological disease, whereas low-dose infection was innocuous, often without evidence of infection (9). Transcriptomic analysis of this fatal infection suggested a typical antiviral response, despite a lack of substantial inflammation in infected tissues (10). Some of the 11 original Trinidad TCRV strains were isolated from what were at the time were called Jamaican fruit bats, but subsequent molecular evidence suggests that Jamaican fruit bats are not found in Trinidad (11). It is likely the bats from which the isolates were made were from the closely-related flat-faced fruit bats, consistent with detection of TCRV in this species in Brazil (3).

A survey of metatranscriptome data of neotropical bats identified TCRV sequences in a Jamaican fruit bat captured in the Dominican Republic in 2014 (TCRV DOM2014) and the full viral genome was confirmed from the original biological material (12). Fortuitously, a small kidney fragment from this bat was dry-frozen in the field and subsequently stored in liquid nitrogen. A homogenate of the fragment was inoculated into Jamaican fruit bats in an attempt to isolate the virus and all became infected. The isolate was then used to examine its virology and host response in a Jamaican fruit bat cell line and in experimental challenge of Jamaican fruit bats.

## RESULTS

*In vivo isolation of TCRV from a kidney homogenate*. A kidney fragment of a TCRV-infected Jamaican fruit bat (12) that had been stored in liquid nitrogen since 2014 was inoculated intranasally and intraperitoneally into six Jamaican fruit bats from a pathogen-free, closed breeding colony (9). On days 3 and 13, three bats were euthanized for necropsy to collect liver, kidneys and spleens that were frozen on dry ice and stored at-80°C. RNA was extracted from the tissues for S segment PCR testing and all were positive by conventional PCR (**Figure 1A**), with spleens showing the greatest viral RNA concentrations by qPCR (**Figure 1B**). The PCR products from some of these were submitted for Sanger sequencing and they were identical to TCRV DOM2014 (12). Homogenates of the day 13 livers were generated, centrifuged to pellet debris, filtered, aliquoted and frozen to prepare a virus stock.

**Figure 1.**
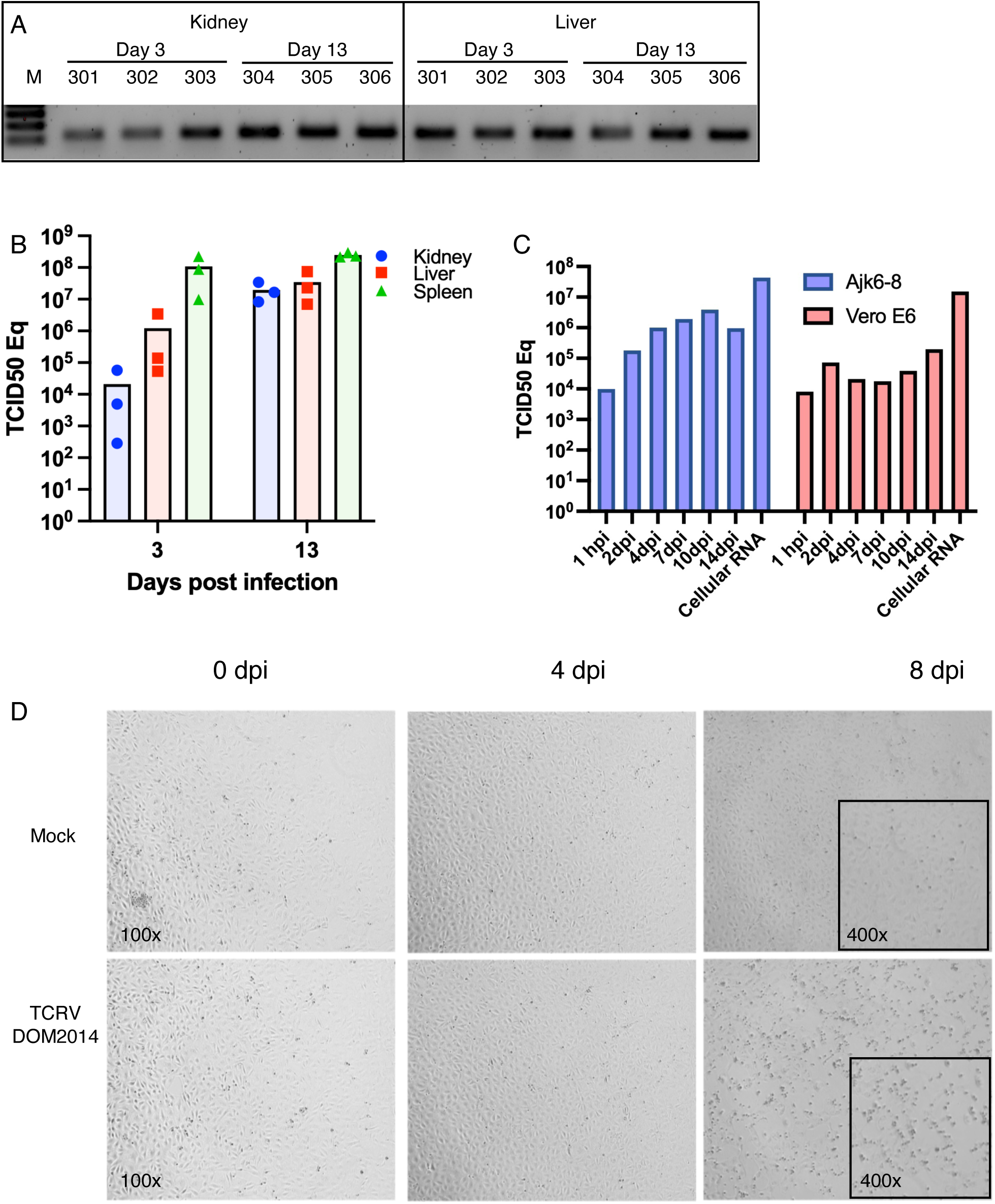
Isolation of Tacaribe virus from experimentally-inoculated Jamaican fruit bats. (A) Six Jamaican fruit bats inoculated with a kidney homogenate of a TCRV+ Jamaican fruit bat captured in the Dominican Republic were PCR+ in both the kidneys and spleens when euthanized on days 3 or 13. (B) Quantitative PCR of viral loads of TCRV S segment. (C) Quantitative PCR of viral loads in supernatants of either Ajk6-8 or Vero E6 cells on days 0-14 and from cellular RNA extracted on day 14. (D) Cytopathic effect of TCRV DOM2014 on Ajk6-8 cells.

*Jamaican fruit bat kidney cell line Ajk6-8 is permissive for TCRV DOM2014*. We have generated several Jamaican fruit bat primary kidney cell lines (13, 14), including several clones produced by flow cytometric sorting. One clone, Ajk6-8, expresses Tfr1 that is used by New World arenaviruses, including TCRV, and this clone was tested for susceptibility to TCRV DOM2014. The line supported virus replication with vRNA levels greater than on Vero E6, although similar amounts of vRNA were detected in cellular fractions (**Figure 1C**). Subtle CPE became apparent on day 6 that became more pronounced on day 8 (**Figure 1D**), demonstrating its susceptibility and permissiveness to the virus. Ajk6-8 cells were used to generate two passages of stock virus for subsequent studies. Genome sequencing of the passage 2 (p2) stock was identical to the original TCRV DOM2014 (12).

*TCRV DOM2014 causes mild pathology in Jamaican fruit bats*. Five Jamaican fruit bats were inoculated with 4×10^5^ p2 TCRV oronasally and intraperitoneally, and were euthanized 10 days later. Although the bats did not exhibit significant signs of disease, some of them had mild jaundice and pale livers, and serum chemistry suggested liver dysfunction, with decreased albumin and blood-urea-nitrogen (BUN) levels (**Figure 2A**). Livers, kidneys, spleens and brains from all 5 bats were qPCR+ for virus, with the greatest vRNA loads in the livers and spleens (**Figure 2B**). Brain was also qPCR+ on day 10 but at lower levels. Three naive bats co-housed three days after infection with inoculated bats were all infected 11 days later (day 13 of the study), demonstrating contact transmission. Viral RNA was still detected on day 21 when the study was terminated.

**Figure 2.**
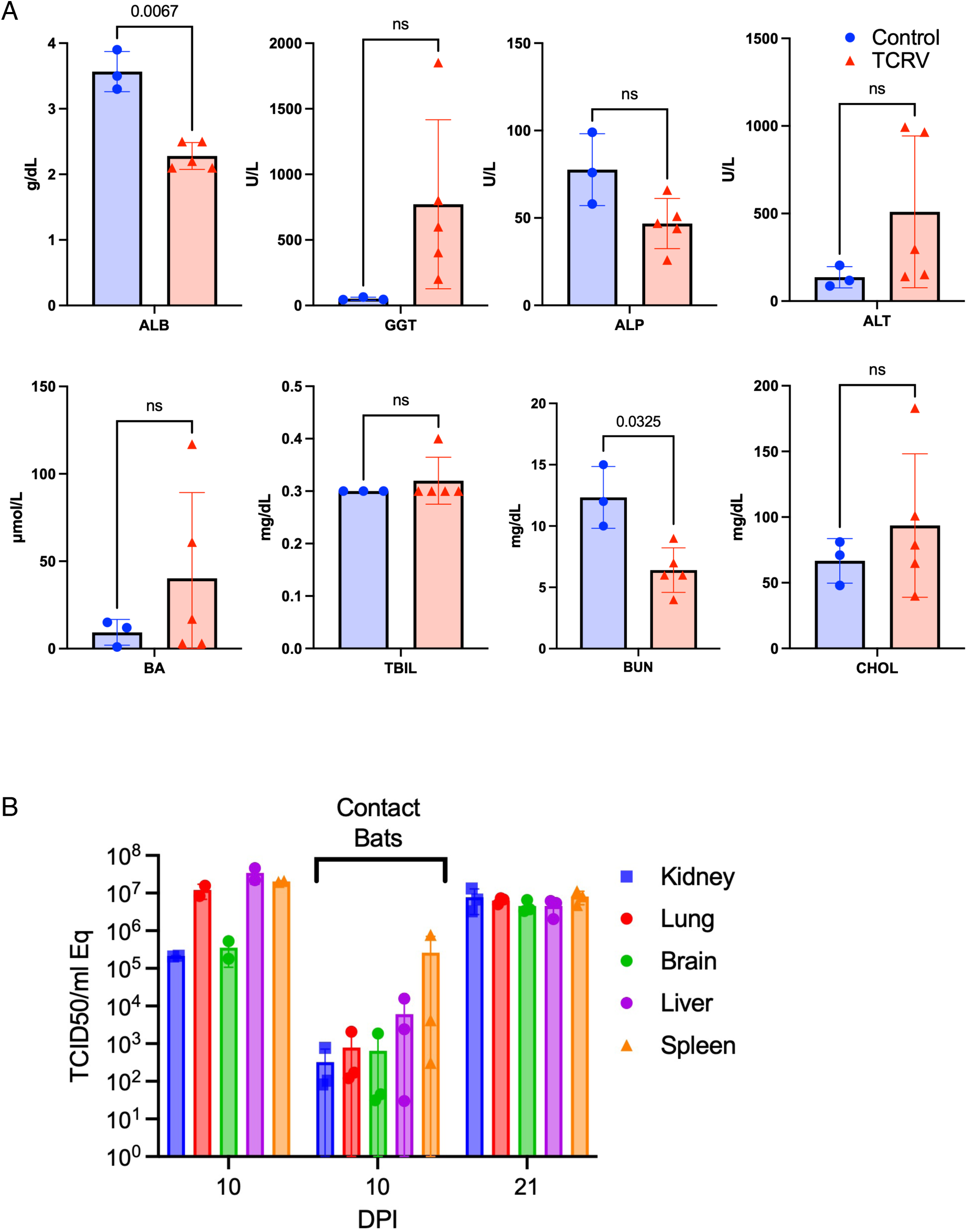
Serum chemistry, viral loads and contact transmission in TCRV-infected bats. (A) Five infected and 3 uninfected Jamaican fruit bats exhibited serum chemistry profiles of liver disease with decreased albumin and blood-urea-nitrogen levels. (B) Quantitative PCR analysis showed infection in multiple organs that increased between days 10 and 21, suggestive of persistent infection. Three contact bats exposed to bats inoculated two days prior all became infected but at lower levels of vRNA detection.

Although livers appeared grossly normal, all bats exhibited mild liver pathology upon histopathological examination (**Figure 3**), including multifocal hepatocyte necrosis with minimal to mild mononuclear inflammation and mild biliary reduplication. Hepatocytes had prominent antigen labeling of glycoprotein by IHC; however, some biliary epithelial and mononuclear cells also labeled strongly for antigen consistent with infection. Two infected bats exhibited mild multifocal tubular epithelial necrosis; however, one uninfected bat was similarly affected indicating that this may be a nominal feature of colonized Jamaican fruit bats. One infected bat had necrotizing and neutrophilic adrenalitis. All infected bats had heavy viral antigen labeling in spleens. Due to non-specific labeling of renal tubule cells in control bats, detection of virus in kidneys by IHC was not possible. Low to moderate levels of viral antigen were present in mononuclear cells of the spleens and lungs as well as neurons and Purkinje cells in the brain, with an increased brain labeling at 21 days post infection as compared to day 10 days post infection. The bat with adrenalitis had positive viral antigen labeling. None of the bats produced neutralizing antibody (titers <20, not shown).

**Figure 3.**
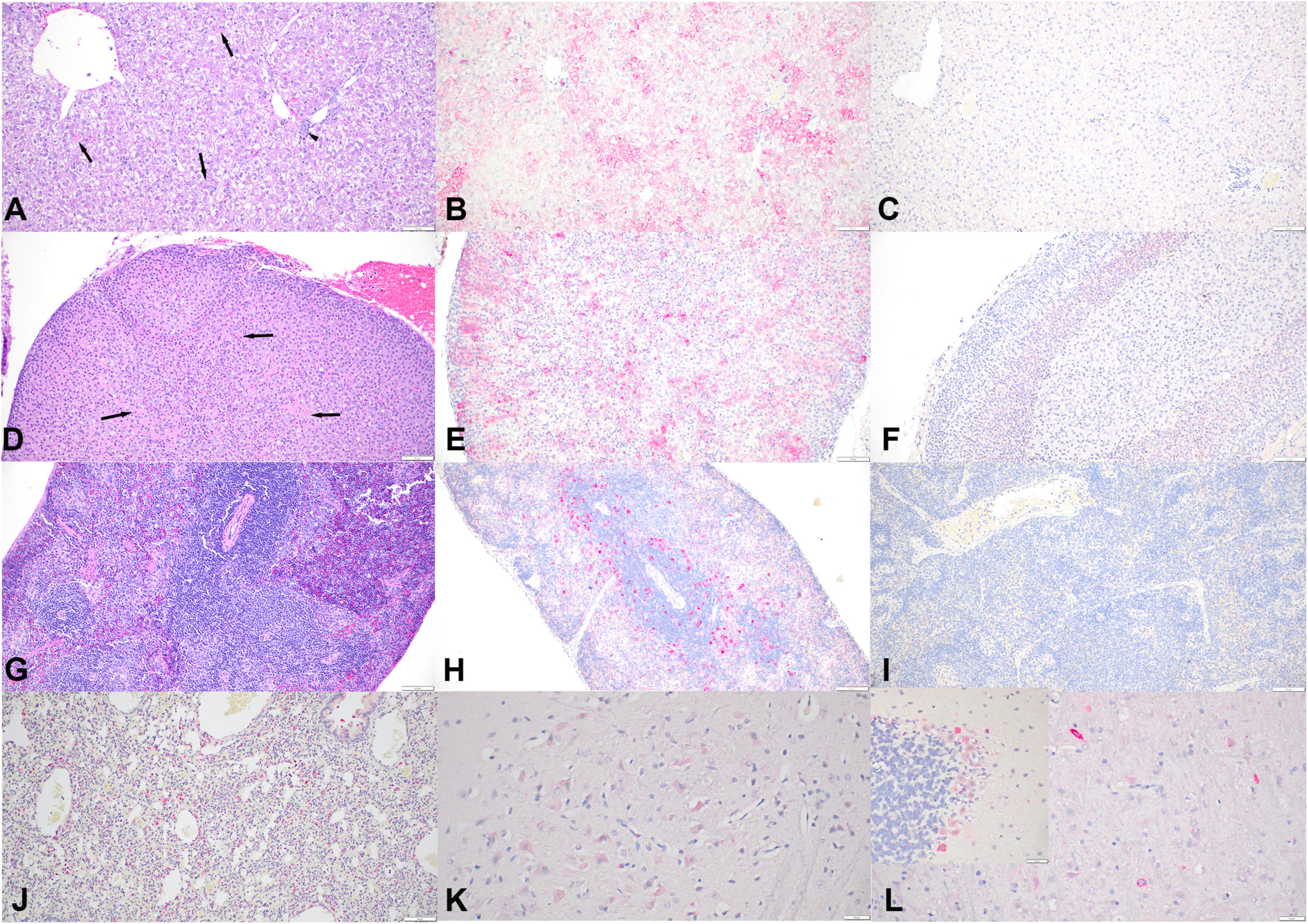
**Histologic lesions and immunohistochemistry**. Infected bats showed individual hepatic necrosis (A, arrows) with mild portal infiltration by lymphocytes and macrophages (A, arrowhead) and multifocal adrenal cortical necrosis (D, arrows). The spleens (G) were histologically within normal limits. Immunohistochemistry of infected bats showed strong cytoplasmic immunoreactivity of hepatocytes (B), adrenal gland epithelial cells (E), mononuclear cells in the spleen (H), macrophages in the lungs (J), and brain (mild labeling of neurons at day 10 post infection (K) and strong labeling of neurons (L) and Purkinje cells (insert) at 21 days post infection). In uninfected bats, there was no immunoreactivity in the liver (C), adrenal gland (F), and spleen (I). Bars = 50 µm (A-L) and 20 µm (L insert). A, D, G: Hematoxylin and eosin. B, C, E, F, H, I, J, K, and L: anti-arenavirus glycoprotein antibody.

*Host response transcriptome profiles.* Because of the high viral loads in livers and spleens, we performed bulk RNA sequencing on these organs from the 5 infected bats 10 days after infection, and 3 uninfected bats. Nearly complete S and L segments were found in all 5 infected bats. KEGG pathway analysis revealed more than one hundred differentially-expressed pathways in each organ. Although many pathways were modulated, immune response pathways elevated in the spleens suggested a typical innate antiviral response, including those associated with viral sensing, type I interferon activation, RNA degradation, ubiquitin activation and MHC class I antigen processing (**Figure 4**). Additionally, autophagy and DNA damage and repair pathways were upregulated, consistent with repair of tissue damage (**Figure 3**). In contrast, adaptive immune response pathways and some innate pathways were substantially repressed, including B cell receptor and T cell receptor signaling, T helper cell differentiation, phagocytic, endocytic, JAK-STAT and mTOR signaling pathways. Inflammatory pathways were also repressed, including NFκB, TNF and chemokine signaling. In addition, many extracellular matrix protein transcripts were repressed, including those involved with cell-cell adherence and leucocyte endothelial migration. For some pathways, certain transcripts were upregulated, whereas others were downregulated.

**Figure 4.**
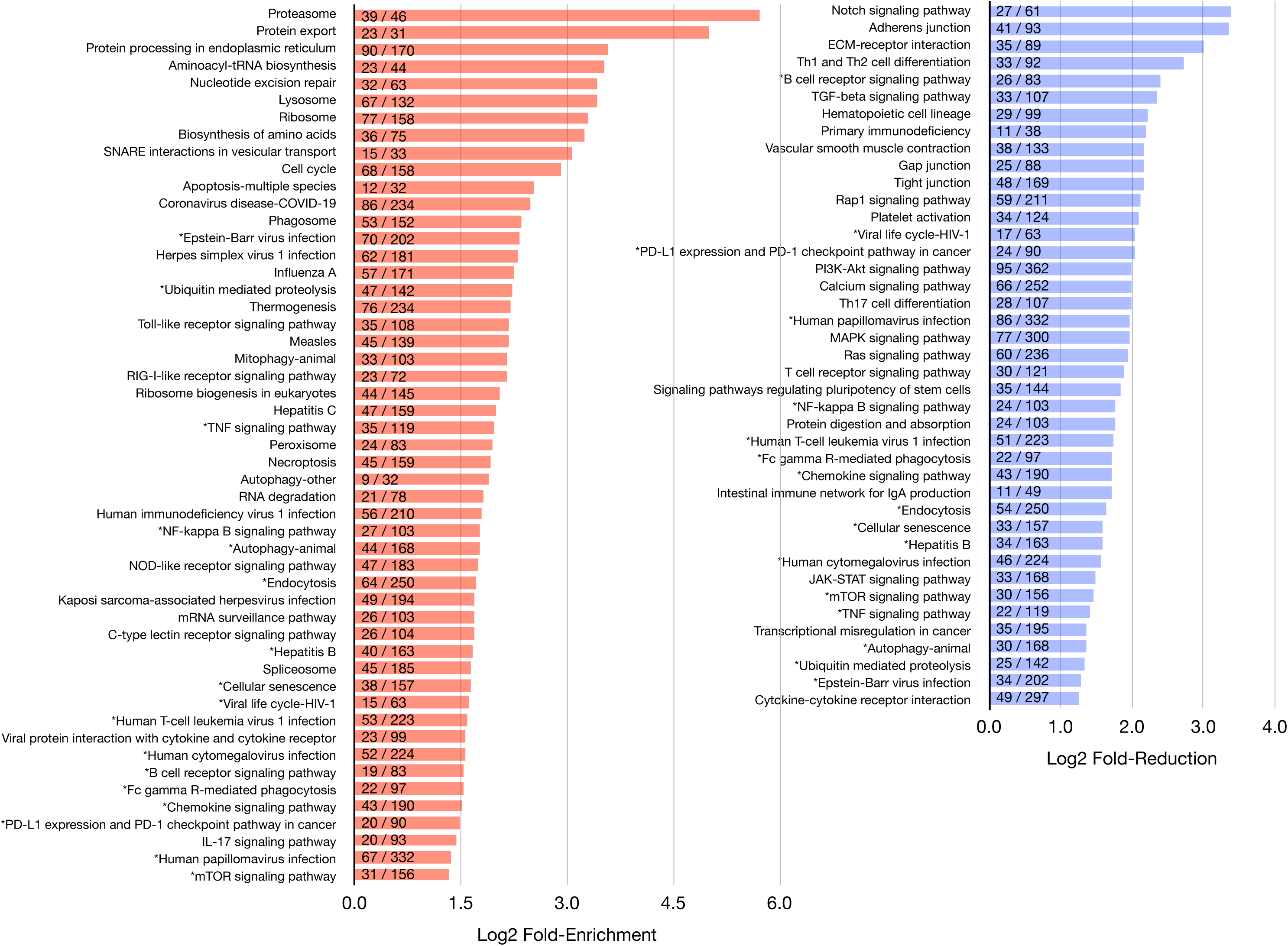
**Immune gene KEGG analysis of spleens from Tacaribe virus-infected Jamaican fruit bats**. KEGG analysis of immune pathways of the spleen showed a typical of an antiviral innate immune response but a repression of pathways associated with B cell and T cell responses. Genes from some pathways were upregulated while others were downregulated are denoted with ‘*’.

Differentially-expressed immune pathways of the livers were fewer but with some overlap with those in the spleens (**Figure 5**). Upregulated pathways were largely consistent with the spleens and suggested a typical antiviral response. However, several antiviral pathways that were upregulated in the spleens were repressed in the livers, including proteosome, peroxisome, lysosome and protein export pathways. Th17 differentiation pathway was elevated in livers but not in spleens.

**Figure 5.**
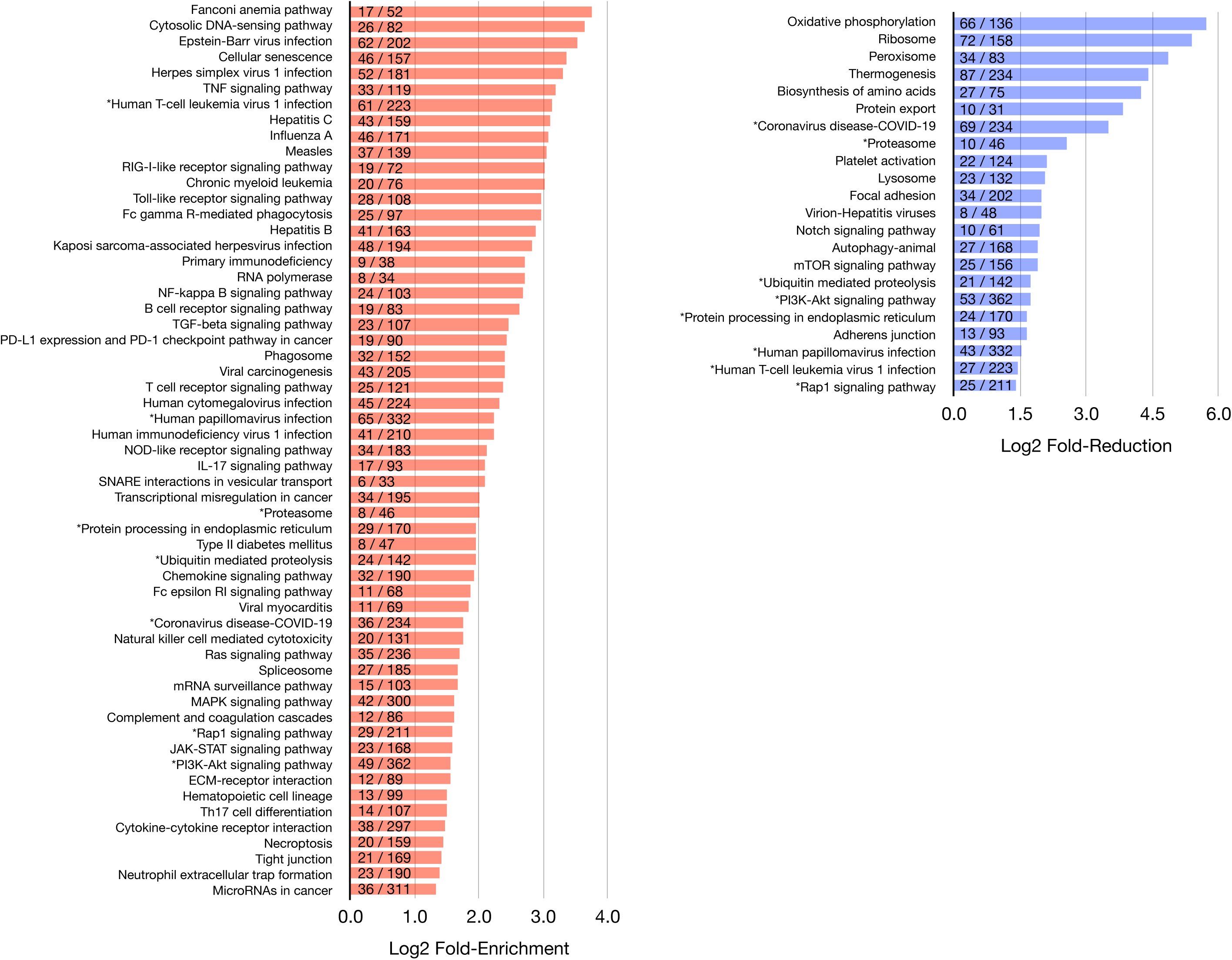
**Immune gene KEGG analysis of livers from Tacaribe virus-infected Jamaican fruit bats**. KEGG analysis of immune pathways of the liver showed a typical of an antiviral innate immune response but some pathways were repressed. Genes from some pathways were upregulated while others were downregulated are denoted with ‘*’.

*Adaptive immune responses to TCRV may be T cell independent.* To assess the impact of TCRV infection on the adaptive immune response, we extracted B cell receptor (BCR) and T cell receptor (TCR) repertoires from bulk RNA-seq data with MiXCR (15). Sequences associated with all BCR and TCR isotypes were identified in the dataset; however, only BCR sequences for IGHM (Igµ), IGHG (Igγ), IGL (Igλ), TRA (TCRα) and TRB (TCRβ) sequences were used for downstream analysis due to low sequence numbers all other isotypes (**Supplemental Table 1**). We assessed basic characteristics of the BCR and TCR repertoires, including V gene usage, somatic hypermutation (SHM) frequency, complementary determining region 3 (CDR3) length, and clonal diversity.

Generally, BCR variable (IGHV) gene usage was similar between infected and uninfected bats with no statistically significant differences in usage between the treatment groups (**Figure 6A**). As in another infection study (16), repertoires of both uninfected and infected bats were enriched for IGHV3 and IGHV4 usage relative to other V segments (**Figure 6A**). However, no statistically significant differences in gene usage were observed between infected and uninfected bats for any individual V gene (**Figure 6A**). Additionally, no statistically significant differences were observed in usage of any individual IGL, TRA, or TRB V gene. However, in an examination of community composition and similarity as a function of clonal identity, infection status differentiated repertoires in both the IGHM and IGHG compartments, suggesting that infection may drive global changes in these repertoires that are not detectable as differential expression of individual V genes (**Figure 6B**; Mantel test: IGHM: r = 0.561, p = 0.019; IGHG: r = 0.639, p = 0.025). In IGL, TRA, and TRB sequences, infection did not differentiate these repertoires (Mantel test: IGL: r = 0.18, p = 0.2; TRB: r =-0.14, p = 0.7; TRA: r =-0.14, p = 0.8), suggesting infection does not strongly influence the IGL or TCR compartments (**Figure 6B**).

**Figure 6.**
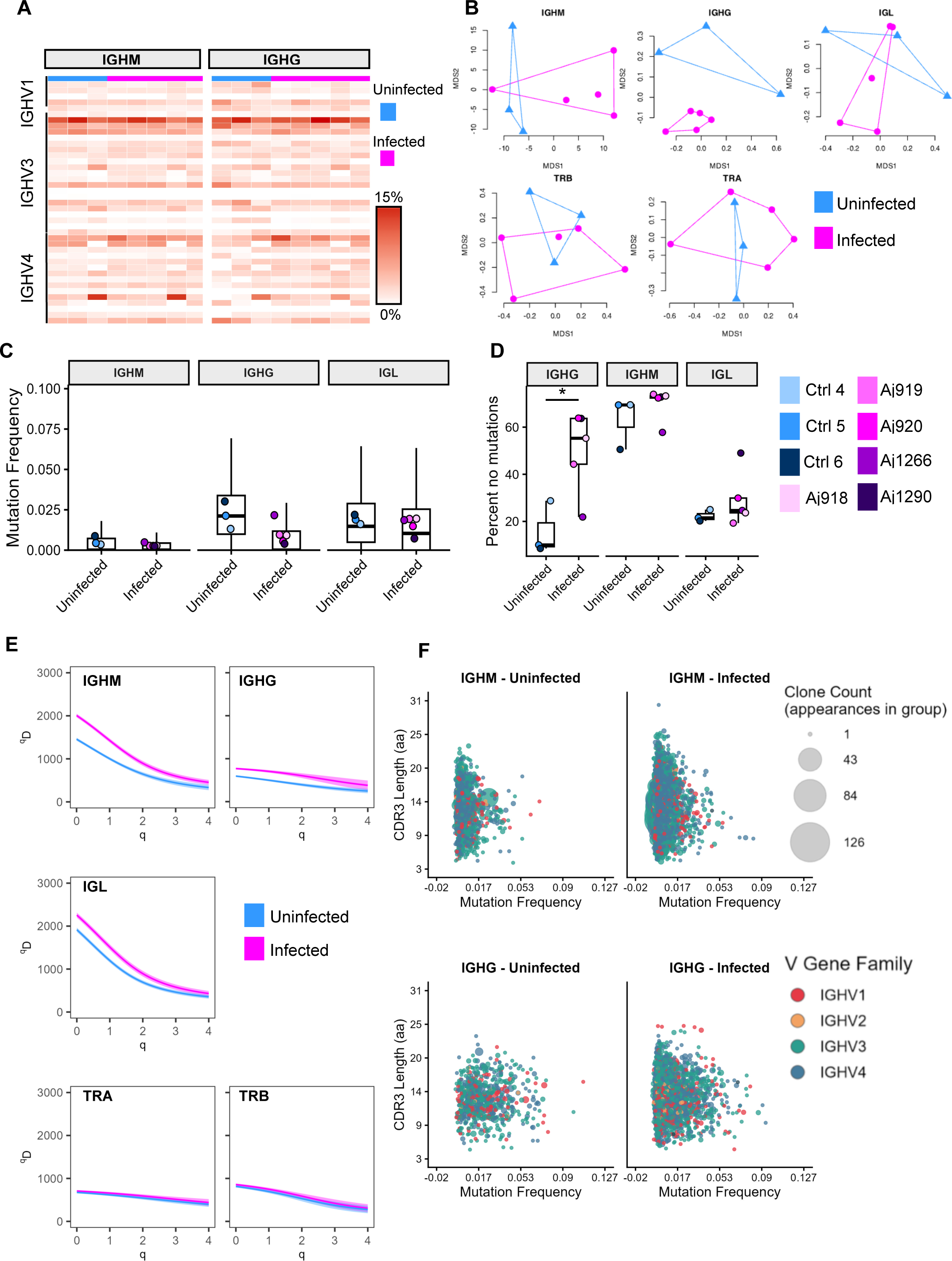
**BCR and TCR repertoire analysis**. (A) Heatmap of IGHV gene usage in IGHM (left) and IGHG (right) repertoires in uninfected (blue; n=3) and infected (pink; n=5) bats. (B) Nonmetric multidimensional scaling plots of uninfected (n=3; blue) and infected (n=5; pink) repertoires based on clonal distribution based on Chao dissimilarity matrices (except IGHM, which is based on the Robust Aitchison distance). Dots represent individual bats and lines border the minimum convex polygons. (C) Box plots of replacement mutation frequency in IGHM, IGHG, and IGL sequences across the entire V region in uninfected (n=3) and infected (n=5) bats. No differences were statistically significant as determined by two-way mixed measures ANOVA with pairwise comparisons using Tukey’s test. (D) Box plots depicting the percentage of IGHM, IGHG, and IGL sequences with no replacement mutations across the entire V region in uninfected (n=3) and infected (n=5) bats. * p < 0.05 as determined by two-way mixed measures ANOVA with pairwise comparisons using Tukey’s test. (E) Diversity of IGHM, IGHG, IGL, TRA, and TRB repertoires for uninfected (n=3) and infected (n=5) bats. For this analysis, all individual repertoires were combined based on isotype and diversity analysis was performed for each combined isotype dataset. In each plot, higher qD (also known as Hill number) values indicate greater diversity within the sample, with q denoting the order of diversity. Each line represents all samples analyzed within an isotype and the ribbon indicates the 95% confidence interval as determined by bootstrapping. Overlapping ribbons indicate no significant difference between repertoires. (F) Bubble graphs depicting clonal diversity and characteristics of IGHM and IGHG repertoires from uninfected (n=3) and infected (n=5) bats. Each bubble represents a unique clone, and bubble sizes were determined by the number of sequences belonging to each clone. Bubbles are colored based on the IGHV gene family associated with each clone and plotted based on the length of the CDR3 sequence associated with the clone and the average replacement mutation frequency across the entire V region of all sequences associated with the clone. In all box plots, the center line indicates the median, upper and lower limits of the box indicate the first and third quartiles, and the whiskers indicate the upper and lower limits of the data within 1.5 times the interquartile range. Dots indicate average values of each individual sample.

Across IGHM and IGHG repertoires, lower SHM occurred in infected bats compared to control bats (**Figure 6C**). We also observed significantly more IGHG sequences with no replacement SHM in infected bats than control bats (**Figure 6D**).

Overall diversity was increased in IGHM, IGHG, and IGL repertoires in infected bats compared to uninfected bats (**Figure 6E**). Across individuals and infection types, IGHM repertoires were more diverse than IGHG repertoires, likely reflecting a broader IGHM compartment composed of naive B cells and an IGHG compartment shaped by selection pressure from previously encountered pathogens. No differences in TRA or TRB diversity were observed (**Figure 6E**). In infected bats, both the IGHM and IGHG repertoires were composed of many clones, each supported by only a few sequences, suggesting that infection does not drive the expansion of selected clones, but rather an increase in the number of unique clones (**Figure 6F**). In infected bats, these clones generally had low levels of SHM and CDR3 lengths that were comparable to uninfected bats (**Figure 6F**).

*Metabolic pathway transcriptome profiles.* Eighty metabolic pathways were represented in a KEGG pathway analysis, and 68 unique metabolic pathways were identified as significantly (Enrichment FDR < 0.05) regulated by TCRV infection within the liver, spleen or both (**Figure 7**). Of the 68 significantly dysregulated metabolic pathways, 55 were downregulated in the liver, 4 were upregulated in the liver, 5 were downregulated in the spleen and 39 were upregulated in the spleen (see **Figure 7A**, set size).

**Figure 7.**
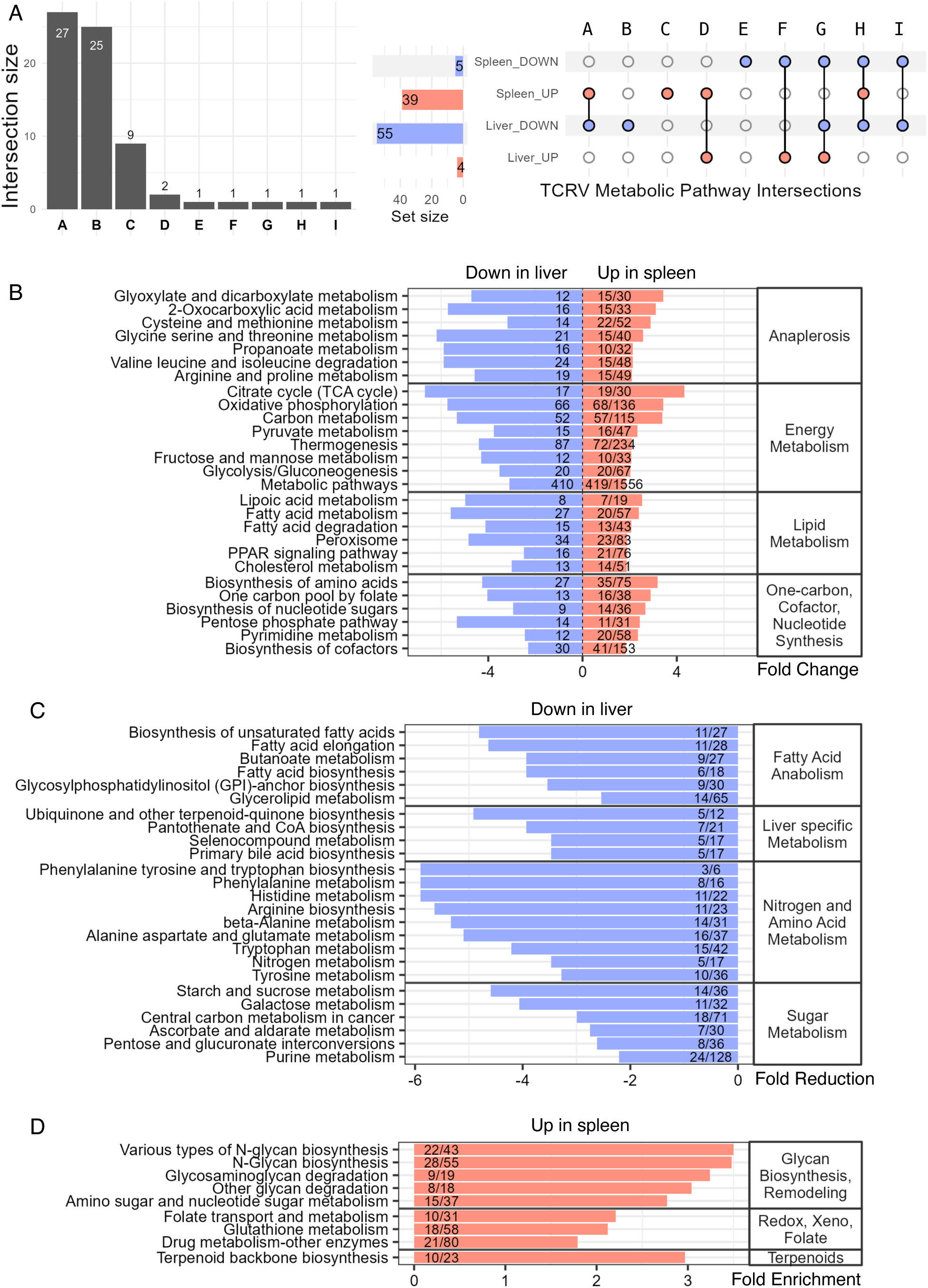
Metabolic transcripts are disparately dysregulated in the livers and spleens of infected bats. (A) Upset plot describing the intersection of significantly dysregulated (Enrichment FDR < 0.05) metabolic pathways between liver and spleen. Left; The intersection size (gray bars) describes the number of pathways commonly dysregulated between the indicated experimental conditions (intersections A through I). Right; The colored dots under each intersection label indicate which tissue-direction intersection the respective gray intersection bar describes. The set size describes the number of significantly up-(red bars) or downregulated (blue bars) pathways in each tissue. The largest intersection was observed for 27 metabolic pathways that were significantly upregulated in the spleen and significantly downregulated in the liver. Twenty-five metabolic pathways were uniquely downregulated in the liver, and 9 pathways were uniquely upregulated in the spleen, none of which were significantly different in any other experimental condition when compared to mock. (B) Fold change of individual pathways (left axis) that were significantly elevated in spleen and downregulated in liver (intersection A). Pathways were grouped by function (right axis). (C) Pathways downregulated in the liver (intersection B). (D) Pathways significantly upregulated in the spleen (intersection C). The numbers within each bar for plots B-D represent gene hits / total pathway genes.

Twenty-seven pathways were both significantly downregulated in the liver and upregulated in the spleen (**Figure 7B**, intersection A), suggesting a disparate or opposing metabolic phenotype between the two tissues. Multiple pathways link to central carbon metabolism, anaplerotic sources that feed the TCA cycle and nucleotide biosynthesis, which may corroborate transcript upregulation to fulfill metabolic requirements for immune cell activation and proliferation in the spleen.

Twenty-five pathways were uniquely downregulated in the liver (**Figure 7C**, intersection B), 9 of which were related to hepatic nitrogen and amino acid catabolism that are major functions of healthy livers. Other groups of downregulated pathways were related to hepatic sugar metabolism and fatty acid and lipid biosynthesis. Highly liver specific bile acid synthesis, pantothenate and CoA biosynthesis, ubiquinone and selenocompound biosynthesis pathways were also downregulated. Nine pathways were uniquely upregulated in the spleen (**Figure 7D**, intersection C). These functions do not have direct liver counterparts. Upregulated in both the spleen and liver (**Figure 7A**, intersection D) were nucleotide metabolism and lipid and atherosclerosis, suggesting elevated synthesis of cholesterol. Downregulated in both spleen and liver (intersection I) was lysine degradation. Upregulated in the liver and downregulated in the spleen (intersection F) was choline metabolism in cancer. Uniquely downregulated in the spleen (intersection E) was inositol phosphate metabolism. Proteoglycans in cancer (intersection G) and glycerophospholipid metabolism (intersection H) were also found to be significantly both up and down regulated during infection.

## DISCUSSION

Many NW arenaviruses that infect humans are BSL-4 agents and model arenaviruses that require lower biosafety containment are needed to study reservoir host infections (17–19). TCRV is a BSL-2 virus that has been used as one such model (9, 20, 21). We have isolated a new strain of TCRV from a wild-caught Jamaican fruit bat that has only been passaged in Jamaican fruit bats or its cells that likely retains its biological and virological characteristics. The virus causes a robust infection in experimentally-challenged Jamaican fruit bats but without visible signs of disease, despite alterations in serum liver profiles and substantial viral burden in the livers and spleens, and which can be transmitted to contact bats. The virus is readily propagated in a Jamaican fruit bat clonal kidney cell line, Ajk6-8, that we derived by flow cytometric sorting and thus makes this model among the most attractive for conducting bat virological, immunological and metabolic studies.

We determined that Ajk6-8 expresses transferrin receptor-1 used by TCRV and other NW clade B arenaviruses, including those that are human pathogens. This clone yielded a high titer of the virus and was used to generate cell culture stocks. CPE became visible on day 6 with maximal virus production on day 8. The DOM2014 p2 cell culture stock generated with this cell line retained the sequence fidelity of its original genome (12).

DOM2014 RNA was detected in all four organs examined by PCR; liver, spleen, kidney and brain, as we previously described for TRVL-11573 infection of Jamaican fruit bats. Antigen staining peaked on day 10 in liver but peaked on day 21 in the brain. Infection was consistent with mild liver disease, including slight jaundice and pale livers in some of the bats and changes in liver enzymes suggestive of liver disease. All bats euthanized on day 21 had high levels vRNA suggestive of persistent infection, thus long-term studies will be required to determine whether the virus is cleared. Our previous experimental infection study showed that high dose (10^6^) inoculation of Jamaican fruit bats with TRVL-11573 led to fatal neurological disease starting on day 10 (9) that did not occur with DOM2014 (4×10^5^). It may be that TRVL-11573 has acquired neurological features due to its passage history in mouse brains (1) or that the lower dose used here was insufficient to cause neurological disease. The levels of DOM2014 vRNA were substantially greater in inoculated bats than in the contact transmission bats, suggesting the inoculation dose may have been greater than what occurs in nature. An artificially-high inoculation dose could be the cause of the observed liver pathology. Convalescent antiserum treatment of Junìn virus infections occasionally causes encephalitis in humans (22), suggesting antibody-dependent enhancement. It may be that adoptive transfer of high-titer Jamaican fruit bat antiserum to TCRV could replicate the phenotype observed in humans.

RNA-Seq analysis showed that a typical, robust antiviral innate response occurred in both the spleens and livers. However, signatures of adaptive immunity in both B cells and T cells were repressed in the spleens, suggesting TCRV may ignore the innate response and, instead, target the adaptive response. Previous work in T cell-depleted laboratory mice showed that T cells were responsible for TRVL-11573-induced meningoencephalitis (20), suggesting that viral suppression of T cell response in bats may limit pathogenesis that could favor persistence. In the livers, fewer immune pathways were activated or repressed; however, a small number of transcripts (14 of 107) of Th17 cell differentiation pathway was elevated, as was B cell receptor signaling (19 of 83 genes) that were contrary to what was observed in the spleens. Otherwise, many antiviral pathways were elevated in the livers, again suggesting an expected host response to virus infection.

Analysis of splenic BCR sequences in TCRV DOM2014-infected bats showed no evidence of increased somatic hypermutation (SHM) nor high titer antibody responses. Increased IGHG sequences with no SHM, increased IGHG repertoire diversity and similar IGHG clonal communities in infected bats without a similar change in repertoire diversity of the TCR compartment, suggests the IGHG response may arise through a T cell-independent mechanism but which is insufficient to generate neutralizing antibody, although we cannot exclude protective non-neutralizing antibodies. Infection appears to promote general B cell involvement and broad community change without driving changes in the B or T cell repertoires in specific V gene usage, mutation frequency, or CDR3 length; however, these results come from bulk RNA-seq data, thus granular changes in the BCR or TCR repertoires may be missed. The lack of diversity in TRA or TRB that were observed between infected and uninfected bats suggests TCRV infection does not exert selection pressure on the TRA/TRB compartment. Moreover, it is possible that suppression of the T cell response by the virus may account for the lack of SHM and affinity maturation that normally leads to high-titer serum antibody responses. Rechallenge of Jamaican fruit bats with H18N11 influenza A virus led to significantly higher antibody titers (23), thus it may be that rechallenge studies with TCRV may also lead to neutralizing antibodies. It is also possible that antibody responses are slower to develop to TCRV and that day 21 is too early to detect neutralizing antibodies, similar to hantavirus infection of reservoir rodents (24). A lack or delay of neutralizing antibody could favor virus persistence in bat populations. Future studies examining antigen-specific B cells or different timepoints following TCRV infection may uncover subtle changes in the BCR and TCR response that were missed in our global transcriptome data.

Together, the RNA-Seq and SHM data are consistent with an infection that may not lead to a robust adaptive immune response. Despite this, in all organs viral loads had significantly declined by day 21, other than the brain where it was greater. It may be that the innate antiviral response of Jamaican fruit bats is sufficiently robust that it alleviates pressure on the adaptive response for control of infection.

Our previous Jamaican fruit bat studies with TRVL-11573 failed to elicit neutralizing antibodies although some bats had low titers by ELISA to nucleocapsid antigen (9). Similarly, field samples collected in Trinidad also identified *Artibeus* bats with low antibody titers by ELISA (25). None of the infected bats challenged with DOM2014 generated neutralizing antibodies by day 21, either. Typically, infections lead to an increase in SHM in the IGHG repertoire as class-switch recombination and affinity maturation produce high-affinity, mutated B cells; however, in acute dengue infection in humans, lower SHM rates have been observed in IGHG B cells, potentially related to the induction of a germinal center independent response (26). Other studies with bats have found that they frequently do not generate substantial neutralizing antibody titers and this may be due to reduced somatic hypermutation that leads to affinity maturation, which has recently been shown to be limited in Jamaican fruit bat B cells (16, 27). This observation appears to be associated with viral infection; immunization with inert antigen leads to robust antibody titers in Jamaican fruit bats (27). Thus Jamaican fruit bats are capable of producing high-titer antibody responses but seemingly not to viral infections. Whether this occurs in other bat species is unknown. In other experimental infection studies of reservoir host bats with Marburg virus (MARV) or Hendra virus, low antibody titers were also found (28, 29) but ELISA absorbances to MARV and Sosuga virus were detected nearly three weeks after challenge (30, 31). Unfortunately, optical densities cannot determine end-point titers, making direct comparisons difficult. Yet other studies of wild bats have detected neutralizing antibodies, including vampire bats to a morbillivirus and to rabies virus (32, 33). Thus, bat antibody responses to viruses remains unclear; however, it is likely that different virus families my elicit distinct immune responses in reservoir host bat species.

The liver and spleen metabolic activities are coupled, often termed the hepato-splenic axis (34). The transcriptional response following TCRV infection shows a consistent pattern with known metabolic coupling and reciprocal adaptation in this hepato-splenic axis. During TCRV infection, the metabolic patterns imply the spleen may act as a metabolic sink for immune activation, while the liver reduces broad metabolic throughput, despite serving as the canonical metabolic center. In the spleen, many interconnected pathways are turned on together, forming a single, coordinated “immune metabolism” network. These pathways include central carbon metabolism (glycolysis, TCA cycle, pentose phosphate pathway, and anaplerotic reactions), which collectively boost production of ATP (energy), NADPH (reducing power), and nucleotides needed for immune cell growth and division (34). PPAR signaling, lipid metabolism, and peroxisomal activity are also increased, supporting membrane synthesis, lipid-based signaling, and other processes that help immune cells proliferate and communicate (35–38).

Amino acid and one-carbon metabolism are elevated in the spleen, supplying building blocks for proteins, maintaining redox balance, and supporting nitrogen-dependent signaling (39). At the same time, enhanced lipid remodeling, β-oxidation, and membrane turnover facilitate phagocytosis and antigen presentation, while increased sugar and nucleotide metabolism supports the synthesis of glycans and nucleic acids that are critical for immune receptor function and immune cell expansion.

By contrast, the liver shows a broad, coordinated reduction in many of the same metabolic pathways that are activated in the spleen. This pattern is consistent with a shift away from high oxidative and biosynthetic activity toward stress-responsive roles such as acute-phase signaling, detoxification, and altered lipid handling, likely reflecting inflammatory suppression or mild pathology (40, 41). Reduced amino acid, one-carbon, lipid, and nucleotide metabolism suggests that the liver is limiting global biosynthesis and redirecting resources toward maintenance and repair.

In addition, 25 pathways that were not detected in the spleen were downregulated in the liver, reinforcing the idea that infected livers are disengaging from their usual metabolic workload to mount an acute response to infection. Only four pathways were upregulated in the liver (IL-17 signaling, NK cell–mediated cytotoxicity, fluid shear stress and atherosclerosis, and a bladder cancer–related pathway), pointing to a local innate/Th17-type inflammatory response and vascular/stress remodeling, consistent with Tacaribe arenavirus infection and reported liver inflammation and necrosis in TCRV models (42).

This model provides the first natural reservoir host system to study a bat arenavirus *in vivo* and *in vitro*. Pathogen-free Jamaican fruit bats and cell lines are readily available to the research community from the Chiropteran Resource Facility at Colorado State University, its genome is complete and annotated (43), including BCR loci (16), single cell atlases of lymphoid, intestinal, liver and pancreatic tissues (44, 45), methods for propagating macrophages and dendritic cells have been developed (45), and many cytokines are commercially available (KingFisher Biotech). Husbandry practices for Jamaican fruit bats have been developed over several decades (46) and do not require significant infrastructure investment for their housing and breeding. TCRV only requires A/BSL-2 practices, and reverse genetics systems for it are available (47, 48). Jamaican fruit bats are also a natural reservoir for H18N11 influenza A virus (49, 50), and have also been used as a model for other viruses, including Ebola virus, MERS-CoV, rabies virus, and SARS-CoV-2 (14, 32, 51–53). Together, these features make the Jamaican fruit bat the most tractable bat model currently available for the research community that can provide detailed examination of viral infection and host responses.

## MATERIALS AND METHODS

### Isolation of TCRV DOM2014

All studies were approved by the Colorado State University Institutional Animal Care and Use Committee and the Institutional Biosafety Committee. The frozen fragment of kidney was homogenized in 1 ml serum-free DMEM in a microfuge tube containing a stainless steel ball at 24 cycles/second for 1 min (TissueLyser, Qiagen). The tube was placed on ice for 2 min and homogenization repeated. The sample was centrifuged at 4°C for 3 min at 1,400g to pellet debris, then the supernatant was passed through a 0.45 µm syringe filter to remove bacteria or fungi that may have been present. The clarified homogenate was brought to 1 mL in serum-free DMEM then 50 µl was inoculated intranasally and 50 µl inoculated intraperitoneally in each of 6 male Jamaican fruit bats. On days 3 and 13, three bats were euthanized for necropsy and liver, kidney and spleens were frozen at-80°C. Small fragments of each were used for RNA extraction (RNEasy, Qiagen). Conventional PCR products were examined by agarose gel electrophoresis to confirm size, and some were submitted for Sanger sequencing (Integrated DNA Technologies) to confirm amplicons were TCRV. One-step qPCR (Qiagen) was performed to detect TCRV S segment (primers, TCRV 1376 FW – 5‘-TCTTGCACCTCATCGGCTTT-3’, TCRV 1501 RV – 5‘-TGGTGGGCTTTCTCAATGGT-3’, TRCV 1424 probe – 5‘-FAM-AGTGTCCTGCTCCTCACAGA – 3’). Homogenates of livers from the day 13 bats were then prepared in serum-free DMEM, pooled, filtered and aliquoted to provide an initial stock of the virus.

### Propagation of TCRV DOM2014 on Jamaican fruit bat kidney cell line

Jamaican fruit bat primary kidney cell clone Ajk6-8 was used to prepare virus stocks and determination of virus kinetics. Verification of transferrin receptor expression was determined by PCR (FW 5’-GAGGTCTGTCGTGACTCCCT-3’, RV 5’-GCTGAGCAGTAGGTTGAGATGT-3’) and Sanger sequencing. Ajk6-8 cells were inoculated with dilutions of the liver homogenate to determine TCID_50_ titer. The liver homogenate titer was then determined on Ajk6-8 cells. A cell culture-derived stock was then made on Ajk6-8 cells by inoculation with a 0.1 MOI of the liver stock homogenate onto Ajk6-8 cells and harvested 13 days later. A p2 stock was then made on the cells using an MOI of 0.1 of the p1 stock and harvested 7 days later.

### Experimental infection of Jamaican fruit bats

Jamaican fruit bats were sourced from the Colorado State University Chiropteran Resource Facility (9). The colony has been closed since 2006 and is pathogen-free. For each study, bats were inoculated with 4×10^5^ TCID_50_ TCRV DOM2014 in 50 µl oronasally and 100 µl intraperitoneally with a total of 10^5^ TCID_50_ TCRV DOM2014. Control bats were inoculated with sterile PBS. Bats were euthanized by inhalation of 5% isoflurane to effect, followed by thoracotomy. Serum from blood collected via cardiac puncture was subjected to serum chemistry analysis (Abaxis VetScan VS2).

### Serum neutralization test

Sera were diluted 1:10 in 2% FBS DMEM, followed by log_2_ dilution series. Neutralization testing (100 µl diluted serum and 100 µl of DOM2014 at 100 TCID_50_) was incubated for 1 hr at 37°C, then transferred to 96 well plate containing confluent Ajk6-8 cells. Plates were scored for CPE on day 8.

### Histopathology and immunohistochemistry

Tissues from necropsied bats were stored 7 days in 10% buffered formalin prior to paraffin embedding and sectioning. Sections were stained with H&E for histopathology, whereas IHC was performed using anti-arenavirus glycoprotein monoclonal antibody (BEI Resources, clone KL-AV-2A1) and anti-mouse IgG-Fast Red. Tissues were read blind by a board-certified veterinary pathologist (coauthor Vilander).

### Transcriptional analysis

RNA was extracted from the spleens and livers from the 5 infected bats and 3 uninfected controls for RNA sequencing (**Supplemental Table 2**). Poly-A libraries for each were generated and sequenced to 40 million 150 nt paired-end reads each (University of Colorado-Anschutz Sequencing Core) and read files were deposited to NCBI (SRA BioProject ID PRJNA1475725). Reads were trimmed with cutadapt (54) prior to processing for differential expression analysis (DESeq2) (55). Transcripts with adjusted *p*-values of 0.05 were submitted for KEGG analysis (56) using ShinyGO (57) and the Jamaican fruit bat reference genome (GCF_021234435.1) (43).

### Germline Ig and TCR annotations

Germline BCR (IGH and IGL) and TCR loci (TRA, TRB, TRD, and TRG) were annotated as we have previously described (16) with some modifications. Briefly, additional IGH and IGL annotations to those we have reported (16) were identified with digger (v0.7.4), a program for *de novo* BCR and TCR germline sequence annotation (58). A reference database composed of IMGT annotations (59) was compiled and position weight matrices from these existing IMGT annotations were created using digger’s internal utilities as described in the documentation (58). TCR annotations were generated with digger; however, TRDD and TRBD genes were located as described in Reers et al. (16). TCR constant genes were identified by aligning TCR exon regions from IMGT (59) to the genome contig containing the TCR V, D, and J gene annotations identified by digger in Geneious (2025.2.2). Regions with IMGT sequences aligned were adjusted for splice sites, when necessary, and translated to check for stop codons. Annotations were combined into fasta files containing either IGH and IGL annotations or TCR annotations for use in BCR and TCR repertoire identification.

### BCR and TCR repertoire identification

Quality of bulk paired-end RNA-seq data was assessed with FastQC (v0.12.1) (reads with a quality score ≥ 30 were retained) and adaptor trimming was performed with Trimmomatic (v0.40) (https://www.bioinformatics.babraham.ac.uk/projects/fastqc) (60). BCR and TCR reads were extracted from bulk RNA-seq data with MiXCR (v4.7.0) (15). Briefly, a custom MiCXR database of Jamaican fruit bat BCR and TCR annotations was built using buildLibrary with-v-gene-feature=VRegion,-j-gene-feature=JRegion,-d-gene-feature=DRegion (for IGH, TRD, and TRB libraries), and-c-gene-feature=CExon1 to create individual reference libraries for each locus. Individual libraries were merged with mergeLibrary and default parameters to create the final reference database. Ig and TCR sequences were identified in the bulk RNA-seq data using MiXCR analyze with our custom Jamaican fruit bat reference library and the arguments --assemble-longest-contigs, --assemble-clonotypes-by CDR3, --keep-non-CDR3-alignments, and --rna. MiXCR results were exported as AIRR files using the command exportAirr and filtered with a custom R script to retain productive sequences with a reported v_call, d_call (for IGH and TRB sequences), j_call, c_call, and junction sequence. Sequences with a primary v_call that was a pseudogene were removed to further filter low-confidence alignments. Filtered files were split based on isotype (IGHG, IGHM, IGHE, IGHA, TRA, TRB, TRD, TRB). Only IGHG, IGHM, IGL, TRA, and TRB files were used for downstream analysis due to low sequence numbers in IGHE, IGHA, TRD, and TRG files.

### BCR and TCR repertoire analyses

SHM frequency in IGH and IGL sequences was determined using the SHaZaM (v1.3.1) module for mutation analysis, available as part of the Immcantation Portal, a collection of Python and R packages for repertoire analysis of BCR and TCR repertoires (61). With the isotype-specific MiXCR AIRR files, the mutation frequency across the entire IGHV or IGLV region for replacement (R) mutations was determined using the regionDefine argument IMGT_V_BY_SEGMENTS.

CDR3 length for IGHM, IGHG, and IGL sequences was calculated using the Alakazam (v1.4.2) module for amino acid physicochemical property analysis (61). Sequence length was determined using default arguments. The first (canonical cysteine) and last (usually tryptophan) codon were removed from the CDR3 sequence prior to analysis.

V gene usage in each BCR and TCR isotype was calculated from the filtered MiXCR AIRR files using *countGenes* with argument *gene=“v_call”* and *mode=“allele”* from Alakazam (v1.4.2) (61).

Diversity of BCR and TCR repertoires identified with MiXCR was calculated using SCOPer and Alakazam (61, 62). While MiXCR assigns clones during the identification of BCR and TCR sequences, it does not assign clones across individuals and cannot account for PCR bias in clonal size estimates from bulk RNA-seq data. Thus, to determine repertoire diversity across infected and uninfected bats, we first combined all sequences from uninfected bats (n=3 individuals) and infected bats (n=5 individuals) into analysis datasets for each isotype (i.e. IGHM, IGHG, IGL, TRA, or TRB). Clones were assigned to sequences with SCOPer (v1.4.0) using the argument *method=“vj”*. After assigning clones, clone size distribution and diversity over a range of diversity orders (q) were calculated using the *alphaDiversity* command from Alakazam (v1.4.2). The 95% confidence interval was determined by bootstrapping (n=100).

When necessary for downstream analysis, clone size was determined by tallying the number of sequences within an isotype dataset assigned to each clone. Mutation frequency of each clone was taken as the average R mutation frequency across the entire V region calculated as described above. To be assigned to the same clonal group, sequences needed to possess the same CDR3 sequence and V gene, so CDR3 length and V gene identity were taken from the first sequence assigned to each clone.

### Community analyses

To assess evidence of convergence between immune repertoires between infected individuals, we used distance matrices to calculate the dissimilarity of the immune “communities”. Clones were defined as described above for each BCR isotype or TCR chain separately. We used the number of each clone found in each individual to calculate a Chao dissimilarity index (63) using the vegdist command in vegan (64) and used Mantel tests (mantel command in vegan) to test whether the community distance matrices and treatment (using a Euclidean distance matrix and dummy variables for treatment) were positively correlated. To visualize the differences in the community we used the metaMDS command in vegan to translate Chao dissimilarity matrices to a two-dimensional representation. In the case of the IGHM clonal communities, there was no good two dimensional nonmetric multidimensional scaling solution based on the Chao index, so these communities are represented visually by an NMDS plot based on the Robust Aitchison (65) distance matrix, but the correlations were still done with the Chao index. Results done with V gene usage instead of clone identity yielded qualitatively similar results.

## Acknowledgements

We thank Mark Stenglein for assistance with RNA-Seq analyses; Jon Flanders, Yolanda Leon, Stephen J. Rossiter, Jackeline Salazar for support during the Dominican Republic 2014.

## Funding

This work was funded by: National Institute of Allergy and Infectious Diseases grants U24AI165424 and R01AI189388 (TS); National Science Foundation IOS 2217296, 2032063, 2031906 and Stony Brook University Presidential Innovation and Excellence Award and NSF-OISE 2020577 (LMD); Experimental Pathology facility at Colorado State University, RRID:SCR_023562 (ACM); and Tracy and Frank Baszler.

## Notes

### Competing Interest Statement

The authors have declared no competing interest.

### Summary of Updates

Duplicate paragraph in description was deleted. A few points of clarification were made to other parts of the manuscript.

